# Properties of Gene Expression and Chromatin Structure with Mechanically Regulated Transcription

**DOI:** 10.1101/262717

**Authors:** Stuart A. Sevier, Herbert Levine

## Abstract

The mechanical properties of transcription have emerged as central elements in our understanding of gene expression. Recent work has been done introducing a simple description of the basic physical elements of transcription where RNA elongation, RNA polymerase (RNAP) rotation and DNA supercoiling are coupled [1]. Here we generalize this framework to accommodate the behavior of many RNAPs operating on multiple genes on a shared piece of DNA. The resulting framework is combined with well-established stochastic processes of transcription resulting in a model which characterizes the impact of the mechanical properties of transcription on gene expression and DNA structure. Transcriptional bursting readily emerges as a common phenomenon with origins in the geometric nature of the genetic system and results in the bounding of gene expression statistics. Properties of a multiple gene system are examined with special attention paid to role that genome composition (gene orientation, size, and intergenic distance) plays in the ability of genes to transcribe. The role of transcription in shaping DNA structure is examined and the possibility of transcription driven domain formation is discussed.

PACS numbers:

## INTRODUCTION

The helical nature of DNA introduces a physical dimension to many important biological processes. One of the most notable is transcription, which is the first step in the conversion of genetic material into biological matter. Though the study of transcription has played a central role in modern molecular biology, much of its physical foundation and behavior is just now being appreciated [2]. Through the use of single-molecule techniques a number of important stochastic and physical aspects of transcription have emerged including the discovery of transcriptional bursting [3–5] and its role in gene expression [6, 7], wide-spread RNA polymerase (RNAP) pausing [8], site-specific transcriptional dependence [9] and the interplay between chromosome structure and function [10]. In this article we will trace the origin of aspects of these phenomena to the physical act of transcription, offering insights into many open problems in biology.

The physical nature of transcription can be conceptualized by the twin-domain model [11] where it was first articulated that transcription and replication cause over-twisting and under-twisting of DNA. The over or under twisting of DNA is referred to as supercoiling (SC) and a number of experimental observations have revealed its central role in transcription [12]. In particular, it can serve as the source of transcriptional bursting [13] and domain formation in bacteria [14].

A recent theoretical framework [1] has turned this decades old conceptual description into a physical model which characterizes the relative amount of RNA polymerase rotation and DNA super-coiling that occurs during RNA elongation. So far, this framework has been applied only to the case of a single RNAP. In this work we will extend this framework so that it can consider the case of multiple RNAPs operating on a common piece of DNA. Additionally, central stochastic elements such as RNAP initiation and mRNA degradation are added to the model. This will allow us to characterize the role of mechanics in gene expression for both isolated as well as interacting genes. The results presented in this work offer plausible, albeit preliminary, explanations for many currently unexplained phenomena in the biology of transcription.

## MATERIALS AND METHODS

### Basic construction of multiple-RNAP multiple-gene system

This article will focus on the characterization of the mechanical properties of transcription and their role in gene expression and DNA organization. The three key physical elements are DNA rotation, RNA polymerase (RNAP) rotation and RNA elongation. Due to the helical nature of DNA, linear RNA elongation is coupled to rotational motion of both RNAP and connected nascent RNA. We will refer to RNAP and nascent RNA collectively as the RNA complex (RNAC). With this in mind, we will need to keep track of both the linear as well as the rotational position of the RNACs as they move along DNA during transcription.

Let us imagine a series of N genes. At a given time there are *M_n_* transcribing polymerases on each gene *n*. The basic coordinates for a given polymerase complex is the distance *x* of the RNAC along the DNA from a transcription start site and the relative rotation of the RNAC *θ* from that site. In the case of multiple RNAC’s on gene *n*, each one has a position 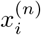 and rotation coordinate 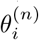. As there are in general many genes present, the start sites are labeled by the coordinate *s_n_* and hence the absolute position of the *ith* RNAC located on the *nth* gene is 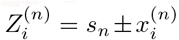; the two signs reflect the two possible gene orientations and thus directions of RNAC movement. A critical variable is the extra DNA twist *ϕ*(*Z*); we will refer to the value of this twist at a particular RNAC location, 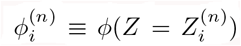, as a linking number constraint (LNC). For an RNAC to move along DNA the topological condition.

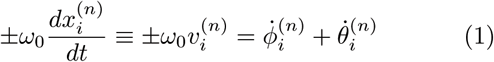

must be obeyed, where *ω*_0_ = 1.85*nm*^−1^ encodes the natural linking number of DNA. Dots denote derivatives with respect to time. Again, the two signs refer to the direction of transcription. The relative difficulty in twisting the DNA (because of opposing torque) or difficulty rotating the RNAC (because of drag) determines the balance between changing *θ* or *ϕ*. For most cases, where the DNA is not completely free to rotate, the relative ratio during transcription can be determined by the balance between DNA torque Ƭ and RNAC drag г(*x*, 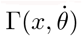) as

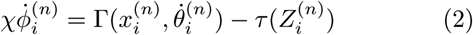

where we have introduced a twisting mobility for the DNA χ and the torque (which is in principle is a functional of the entire *ϕ* field) is evaluated at the RNAC position. As each RNAC moves along DNA its position 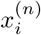 changes, resulting in changes to the dynamic linking number 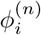 constraint (LNC) at this point.

While many mechanical properties of DNA are well characterized, the mechanical nature of the rotating RNAC is largely unknown. From early studies however it is clear that RNA elongation plays a key role [15, 16]. Even though this is a critical factor, the coefficient and functional dependence of transcript length on the rotational drag Γ are not known at this time. We will posit an RNAC viscous rotational drag which is linear in the rotation speed with a power-law dependence on the transcript length as 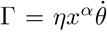 where 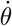 is the angular speed of the RNAC and *η* an unknown coefficient of friction. A length independent drag of the RNAP can be added, however nascent RNA plays the dominant role in generating SC [15, 16] so we will not consider that here. The effects of varying the parameters are examined in the SM and simple convenient choices have been made for the simulations conducted in this work.

Combining equations 1 and 2 results in a coupled dynamic equation for all the different LNCs *ϕ_i_*. First, we can use the RNAC viscous rotational drag term outlined above to derive a dynamic equation for *ϕ*

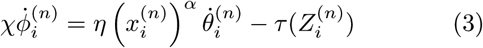

Substituting in the topological constraint eq. 1 we can write

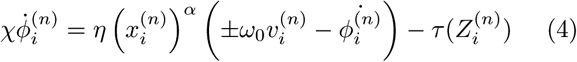

so that we have an equation for the DNA twist in time as

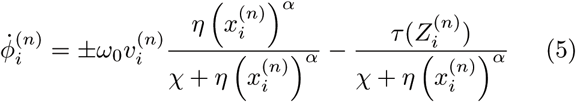

The denominator for both terms is an attenuating factor which incorporates the drag of the RNAP rotation as well as resistance to rotation of the DNA.

Supercoiling and DNA mechanical dynamics occur on a sub-second time-scale [17, 18] whereas typical speeds for transcription are 10 − 50 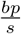 [19]. This means that for genes on the order of 1 *kbp*, transcriptional dynamics happen on the second and minute time-scales. Additionally, RNAP operation is robust against sub-second torque fluctuations [20]. Subsequently, we expect the locally produced supercoiling at the LNC locations to spread throughout the allowed DNA segment on a time-scale faster than that on which transcription occurs. Thus, the torsional response of DNA at the point at which an RNAC is operating 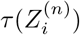 is determined by the state of DNA on either side. More generally, we might expect to solve a supercoiling transport equation with both twisting and writhing degrees of freedom.

With this assumption of complete relaxation, the supercoiling density (SCD) 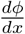 is constant in regions between LNC’s. In particular in front or back of a given RNAC, we have (after normalizing by *ω*_0_)

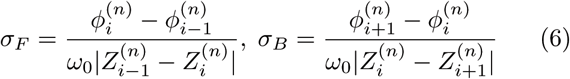

This labeling reflects the fact that the nearest polymerases are the ones with the closest indices. For notational consistency, we must have conditions for when the interaction is between RNAC’s on different genes. Thus, in the case when the neighboring genes to gene *n* as well as gene *n* itself are oriented in the positive direction, we have

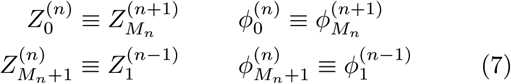

These of course assume that there is at least one RNAC at the neighboring gene; if not we just skip genes until the next RNAC is found. There are analogous expressions (listed for completeness in the SM) for all possible orientations. Finally at the boundaries of the entire system we define M_0_ = M_N+1_ = 1, 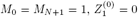, and 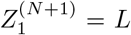. We use a combination of fixed and free boundary conditions which correspond to the introduction of an additional LNC at the leftmost and rightmost edges of the DNA under consideration. The values of *ϕ* at these edges will depend on whether we assume fixed or free boundary conditions.

This above formula shows that the torsional response contains contributions from both in front *τ_F_* of and behind *τ_B_* the RNAC

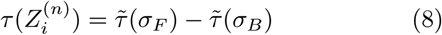

Once these functions are specified, we have a closed from equation for the torsional angles.

Due the previously discussed time-scale separation the torsional (*τ*(*σ*)) response of DNA between two LNCs will be that of steady-state supercoiled DNA. In this framework super-coiled DNA can exist in a purely twisted, purely plectonemic or a mixed state. Following the phenomenological approach given by Marko [21] the torque in a given piece of DNA held at a constant force *f* is specified by the SCD as

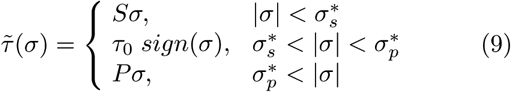

where the coefficients *S,τ_o_,P* and SC transition values 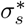, 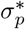 are given by DNA mechanical constants and are a function of applied force (given in [21]). It is worth noting that the introduction of a well-defined applied force is at this time cloudy from an in vivo perspective, its experimental implementation in vitro is straightforward. Additionally, one can interchange the force *f* for a constraint on the average end-to-end distance of the DNA [22].

The simplest example of this system gives rise to an equation for the dynamics of the super-coiling for a simple isolated RNAC operating against a fixed barrier a distance *L* away and with open (zero torque) conditions at *Z* = 0; this was discussed in our previous work. Having a fixed barrier ahead of the the RNAC means that *ϕ* = 0 at *Z* = *L* and having a free rotation behind the elongating polymerase means that 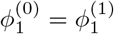 and hence *σ_B_* = 0. To simplify the notation we can use 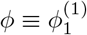 and *σ* = *σ_F_*. In the limit that the gene length is short compared to the barrier distance, the substitution *ϕ* → *σL*ω_0_ yields a dynamic SC equation

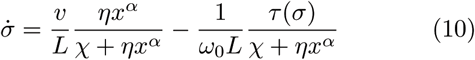

In the limit of very large RNAC drag the DNA relaxation is highly attenuated 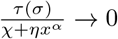 and 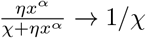 so that 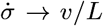. In the opposite limit, the torque injection term dominates so that 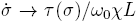. These basic behaviors put bounds on the amount of SC which can be added to a piece of DNA by RNAP during transcription. A more complete examination of this system was recently published [1] and methods for experimentally determining the unknown mechanical parameters of the RNAC drag were outlined.

### Creating a full Model

To make a complete model of the transcriptional process, and hence to allow for the simulation of many RNAC’s on multiple genes, we need to add a few more ingredients. So far, we have not discussed the velocity of the RNAC’s which is obviously needed in the above system of equations. It is well-known that the velocity can depend on the accumulation of SCD in regions between operating dynamic RNACs as well as between RNACs and static boundaries. We will use a simple empirical from of this relationship that qualitatively matches the existent data and accounts for the relatively persistent motion of the polymerase up to some critical value of the opposing torque. Afterwards, we will add in some additional pieces to allow for stochastic transcription initiation, SC relaxation via topoisomerases and RNA decay.

The mechanical properties of RNA polymerase itself are well characterized and it displays constant velocity [19] behavior over a wide range of torque (−20 to +12 pNnm) [20]. Following the previously used RNAP behavior given by [20] we can incorporate a supercoiling dependent velocity by using an expression with logistic-like dependence for each RNAP. The torque dependent velocity observations for SC in front of and behind an RNAP were studied separately; we will consider their simultaneous effects here as

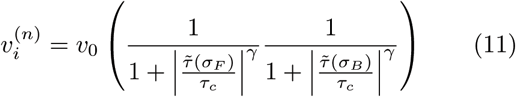

For both negative and positive SC the cutoff torque is near *τ_c_* = 12 (*pNnm*) when applied separately [20]. Though the cutoff for positive and negative SC torque are the same, the severity of the cutoff seems to be more intense for negative SC than for positive but we will ignore that here. Additionally, there are stochastic properties of the stalling that we will not consider here but could be incorporated into future work.

It is clear that the mechanics alone will not stop a trailing RNAP from “passing through” a forward RNAP which is stalled. To prevent passing we have explicitly implemented a hard-core RNAP repulsion so that RNACs cannot come within a distance *δ* of one another. This distance could arise because of a number of physical interactions, however, at this time there is no experimental data to constrain this interaction. We will take the cutoff distance to be the physical displacement caused by the space occupied by RNAP on DNA during transcription. A similar cutoff has been added for the initiation of new RNAP so that another cannot start until the most recent one has moved out of the transcription start site (TSS). Both are fixed at *δ* =15 (*nm*) for all results presented. The of effects of changing the nature and size of the distance *δ* are examined in the SM.

In addition to the mechanical elements outlined above we must include other stochastic elements of transcription to examine the statistical properties of gene expression. We will include stochastic RNAP initiation, decay and topoisomerase action. A new RNAP will start transcribing with a stochastic rate *r* and begin moving from the TSS with a velocity governed by RNAP mechanics given above. A transcript which has been made in its entirety (by an RNAC moving all the way along the length of the gene) will decay stochastically with rate *λ*. Both the initiation and degradation steps are modeled as simple Poissonian processes.

There are generally speaking two types of topoisomerases: topo1 which most effectively removes negative SC and topo2 which primarily removes positive SC [23]. Topo1 breaks a single strand of DNA and allows the DNA to unwind using no ATP while topo2 uses a double strand passing method which uses ATP and is much slower than topo1 [23]. The precise microscopic nature of the relaxation of built up SC, as caused by topoisomerases, is not well known at this time. Consequentially, we use a coarse-grained method for topoisomerase action. For simplicity we remove all the SC between two genes (or between a gene and a static barrier) randomly with rate *g* by replacing the DNA in this region with DNA with SCD that matches the two LNC 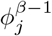, 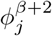 on both sides of the two genes selected for relaxation. This means that if genes *β* and *β* + 1 are chosen for relaxation the LNC and position of the ’end’ RNAC (the RNAC closest or furthest from the TSS depending on the orientation) for genes *β* − 1 and *β* + 2 are used to set the SC level. Explicitly we set

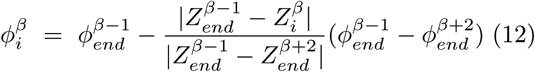

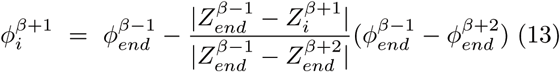

for all *i* ∈ *M_β_, M_β+1_* RNAPs on genes *β*, *β* + 1. For a system with one or two genes this step results in replacing the DNA in the system with complete relaxed DNA with no SC. This is done for two reasons. The first reason is that due to eq.5 the differences in SC between RNACs within a gene tends to be small (see fig.2). The second reason is because the intergenic regions are much larger and homogenous than the regions containing genes for the systems considered in this article. Thus the action of the topoisomerase most like to take place in-between genes. The creation of more sophisticated models of SC relaxation which incorporate more precise modeling of topoisomerase action is left for future work.

**FIG. 1:**
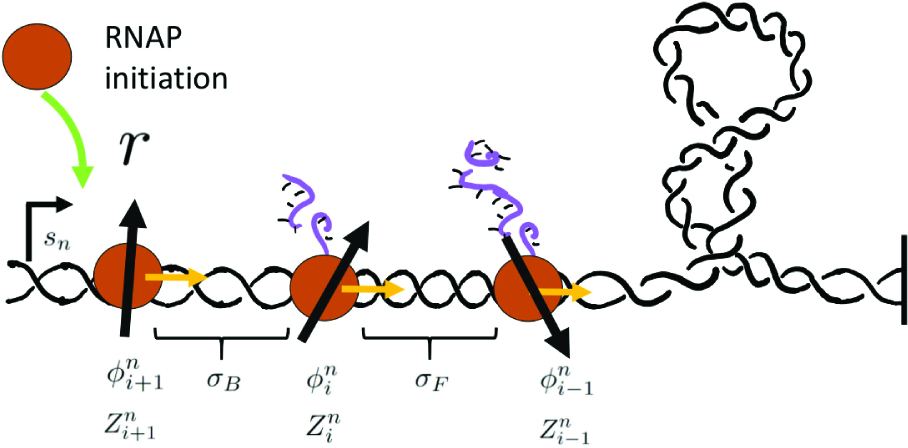
(color online) A cartoon depicting the interaction of RNAPs along a shared piece of DNA. During transcription elongation must occur through a combined rotation of RNAP and DNA. The torsional response of DNA governs the motion of the RNAPs and affects DNA conformations.

**FIG. 2:**
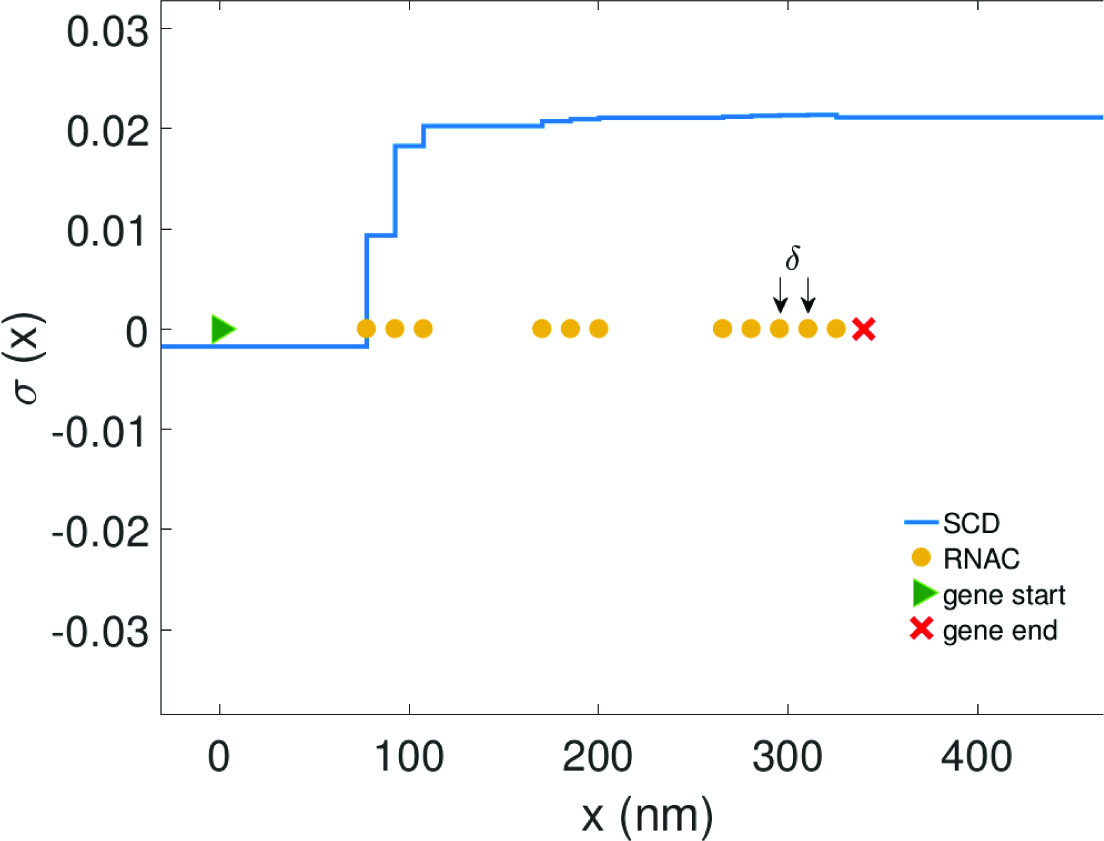
Representative snapshot of RNAC positions in a gene during transcription for an isolated gene torsionally constrained in one direction (composition details in SM). Spacing between RNACs is determined through random initiation, SC dependant velocities and hard core repulsion. Consequentially, as RNACs move along a gene during transcription stalling and clustering occur leading to bursts of mRNA production (fig.3).

## RESULTS

### Results subsection one Bursting statistics for isolated genes

The widespread observations of the stalling of RNAPs and bursting transcription in *in vitro* and *in vivo* systems have been the subject of much research [13, 24–26]. Though much progress has been made, the source and control of these related phenomena is still a matter of debate. In this section we will examine how the mechanical properties of transcription, namely gene and barrier lengths, alter the ability of a gene to transcribe. To do this we will simulate the dynamical motion of multiple RNACs using the framework constructed above for a simple isolated gene with length *GL* with a static LNC a distance *L* from the TSS as shown schematically in fig.. At this time we will not specify the precise nature of the torsional constraint and imagine it as a generic SC barrier. In this configuration positive SC will build up in the region of DNA between active transcription and the barrier.

As positive SC builds the motion of the RNACs is inhibited. Consequentially, RNACs form clusters of various sizes and spacing controlled through random initiation, SC dependant velocities and hard core repulsion. A typical configuration of the system is shown in fig. 2. The clustering of transcribing RNAPs has been observed in a number of experimental studies [27, 28] but a satisfactory mechanism for their formation has not previously been developed. Additionally, this behavior limits the ability of the RNACs to reach the end of a gene and thus affects the production of mature mRNA. Due to this, the production of mature mRNA occurs in burst with periods of active and inactive transcription production emerging in simulations of the system (fig.3). This effect is present only with mechanically regulated transcription (red lines) as opposed to transcription with only stochastic elements (blue lines). Thus, RNAP clustering as well as mRNA bursting robustly emerge from this framework with a common origin in gene composition and mechanics. The size of the bursts vary, because relaxation events can occur between arresting, but happen with a typical value which we will refer to as *m_c_*.

**FIG. 3:**
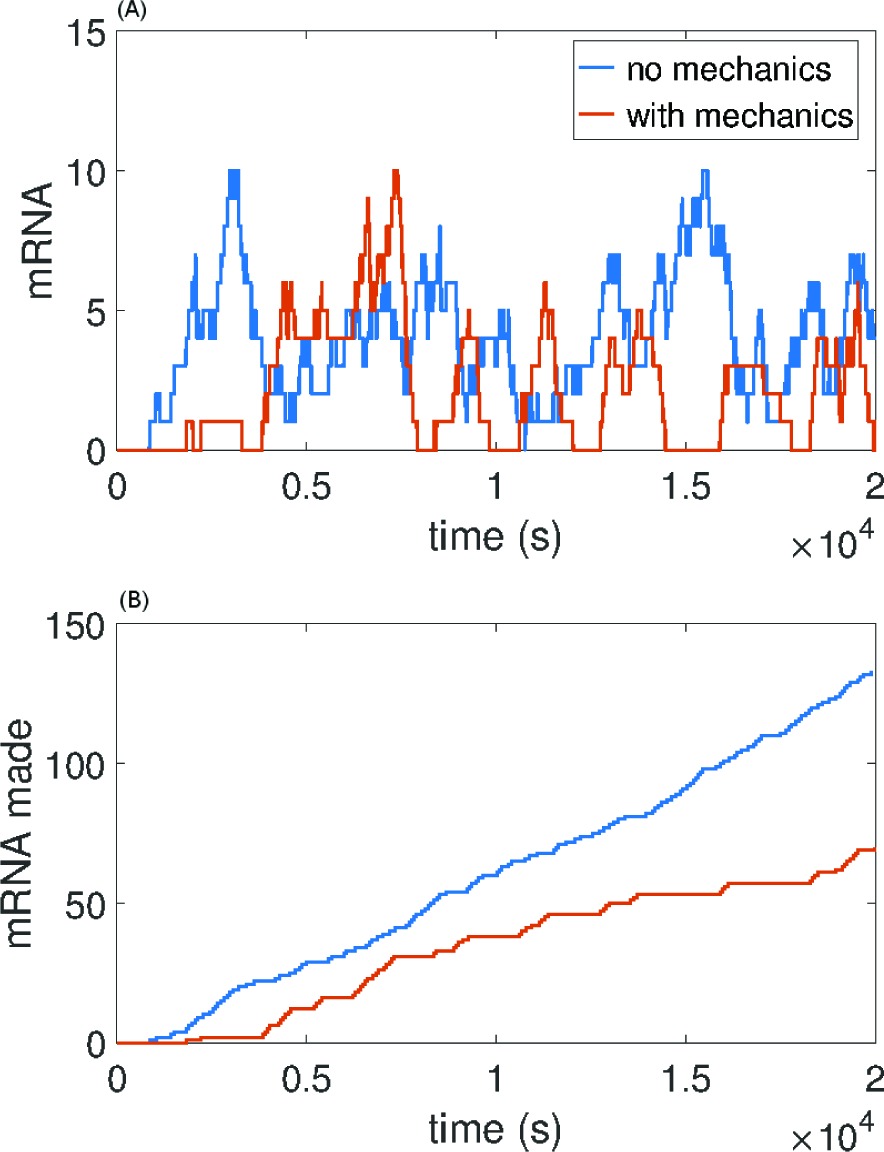
Stochastic trajectories of mRNA present (A) and total mRNA made (B) for an isolated gene torsionally constrained in one direction. Figure (B) clearly shows periods of active and inactive transcription (bursting) for a gene which is mechanically regulated. Gene and barrier lengths are GL=1kbp and L=10 kbp respectively (composition details in SM). Stochastic parameters {*r,λ,g*} = {2^−1^,1/20,1/20(*min*^−1^)} were used. The mechanical parameters {*v_0_,η,α,χ, f*} = {20,1/20,1,10,1} used are held constant for all results unless otherwise stated.

The properties of bursting are controlled by a number of intrinsic and extrinsic parameters that are outlined in the previous sections. However, not all parameters effect the properties of stalling in the same manner. The mechanical form of the drag, through the coefficient of friction *η* and nascent RNA length dependence *α*, influence the quantitative properties of the stalling and burst size *m_c_*. Though their values are unknown at this time, we should imagine that they are fixed for all genes. We shall fix them for all the results presented in the article. Consequentially the role of gene specific parameters, namely gene length (GL) and barrier distance (*L*), as well as external parameters such as force, are of much greater interest as they vary throughout the living world. Unless otherwise specified the values *η* = 1/20, χ = 10, *v*_o_ = 20, *α* = 1 have been used. The effects of changing these intrinsic mechanical parameters are examined in the SM.

We will first examine the role of gene length in bursting. In fig.4A the number of transcripts which are made between stalling (the burst size distribution) is given for three genes with various lengths transcribing against a fixed barrier. Burst size distributions for genes of lengths 500, 1000 and 1500 bps are shown. The characteristic burst size, *m_c_*, emerges as clear mode in the statistical distribution. The data shows a clear relationship between gene length and burst size. As the gene size increases, *m*_c_ decreases and there is also a sharpened burst size distribution. These results arise due to the fact that stalling becomes less frequent for short genes with large *m_c_* because the characteristic time to arrest becomes significantly less than the time between relaxation events. Thus, genes with large *m_c_* remove the competition between stalling and relaxation resulting in Poissonian (constitutive) gene expression. In agreement with this, one can recover a sharper distribution at higher *m_c_* by lowering the relaxation rate; this is shown in fig.4B. Whenever this competition becomes relevant, we obtain regular bursting which can then dictate gene expression statistics, as will be examined later.

**FIG. 4:**
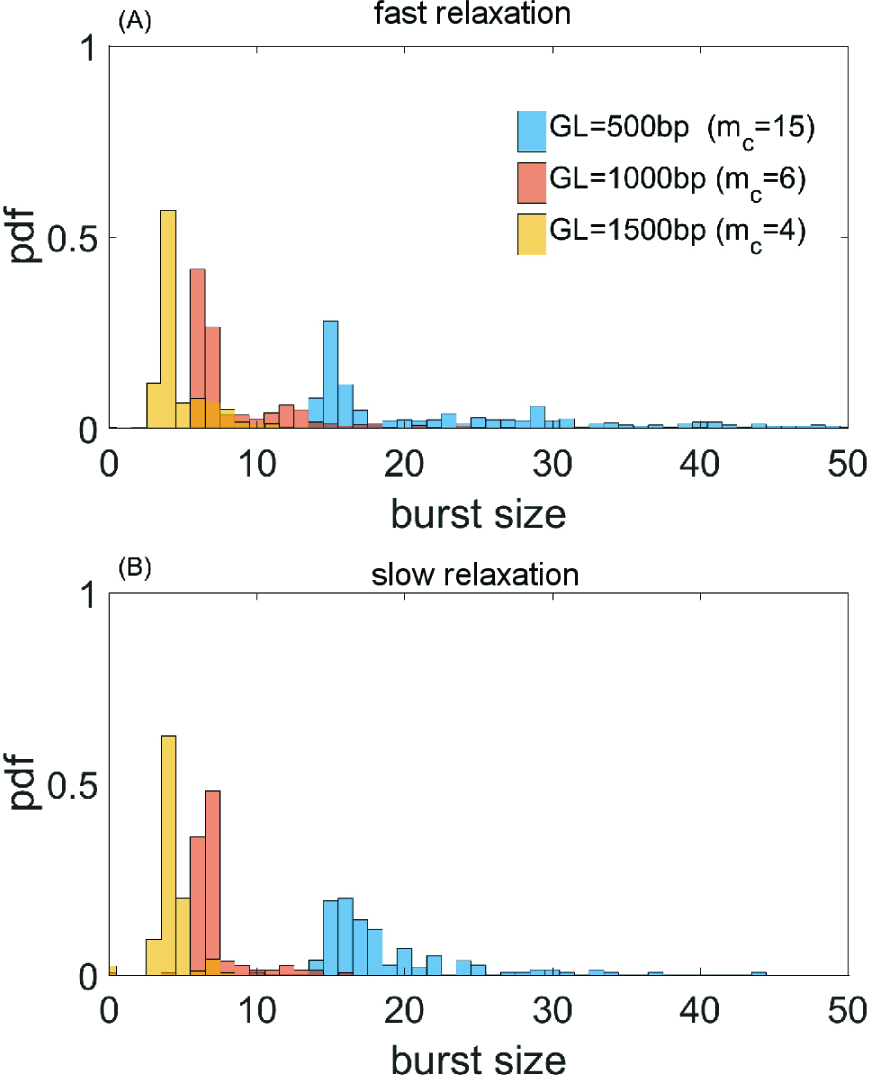
Burst size distribution defined by the number of transcripts made between inactive periods of transcription for three genes of increasing lengths transcribing against a single barrier (composition details in SM). Data is shown for two relaxation rates. A clear mode exists in each distribution corresponding to the typical number of transcripts made between stalling event *m_c_*. As the relaxation rate is increased ((A) *g* = 1/20, (B) *g* = 1/80) the mode is unchanged but the variance in burst size is increased due to decreased stalling frequencies. All other simulation parameters are the same as figure 3.

In addition to changing the gene length the distance to the barrier can be altered. Doing this will also change the burst properties of the gene. The characteristic burst size *m_c_*, as given by the mode of the burst size distributions, is shown in fig.5 for a wide range gene lengths transcribing toward a fixed barrier at various distances. As expected, increasing gene length decreases *m_c_* while increasing the barrier distance increases *m_c_*. As the value of *m_c_* becomes very large (due to short gene length or large barrier distance) bursting becomes infrequent. These properties demonstrate that geometric properties of genes (their length and spacing) play a fundamental role in their operation.

**FIG. 5:**
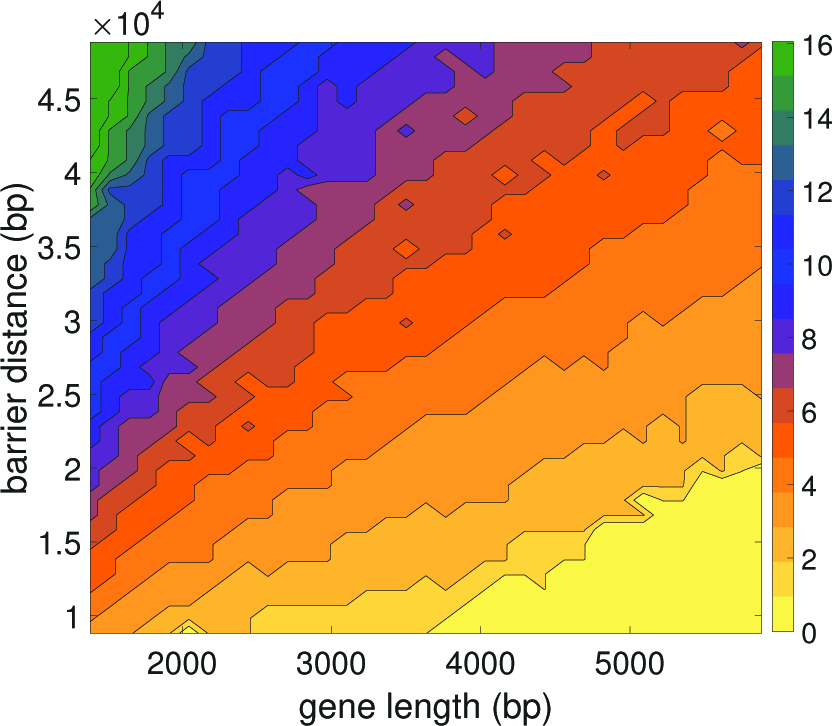
Burst size *m_c_* dependence on gene length and barrier distance for an isolated gene transcribing against a single barrier (composition details in SM). As gene length (GL) decreases the burst size *m_c_* increases and the number of bursts decreases. For very small genes with long barriers (L) the burst size *m_c_* mode is smoothed out do to infrequent bursting (see fig. 4) changing the nature of transcription (see fig. 3).

To get a qualitative handle on how *m_c_* changes as a function of gene and barrier length we can imagine that each transcript introduces Δ*σ* supercoiling during its elongation and that the stalling torque *τ_c_* occurs at a corresponding level of SC, *σ_c_*. In this case the gene can make *m_c_* = *σ_c_/Δσ* transcripts between relaxation events. Writing this more explicitly, the change in SC is given by *Δσ* ~ *ϕ*/*ω_0_W* where *ϕ* is the amount of twist added during the making of a single transcript and *L* is the distance to the barrier. For a long gene, of length *GL*, most of the elongation goes into DNA twist over RNAC rotation so that *ϕ* ~ ω_0_*GL*. Then *m_c_* ~ *L/GL* matching the qualitative behavior of fig. 5. In general, however, an exact analytical calculation of *m_c_* is not possible and will depend on the precise values of the mechanical properties of transcription outlined in the previous section.

The generic changes in the gene expression patterns caused by arrangement of the genes, as well as the difference in susceptibility in gene expression under varying external constraints, might offer an explanation for a number of known gene expression phenomena which are related to changing mechanical properties. These include mechanical transduction of gene expression and cell differentiation [29], diseases related to changes in mechanical properties of the cell [30] as well as direct changes in the constraints and structure of chromatin itself such as been recently observed in cancer [31].

As the external force is increased, genes in the system are affected differentially. This is shown in fig. 6 where short genes undergo small changes as the force is increased while long genes undergo large changes. Since one can interchange the force constraint *f* for a constraint on the average end-to-end distance of the DNA [22] the mechanical consequences of the external constraints as well as of their positions can be examined. The model and results presented in this article are at present too crude to quantitatively examine any specific examples of this type of phenomena. However, the results clearly demonstrate the ability of external physical and mechanical changes to a cell and its DNA to alter its gene expression.

**FIG. 6:**
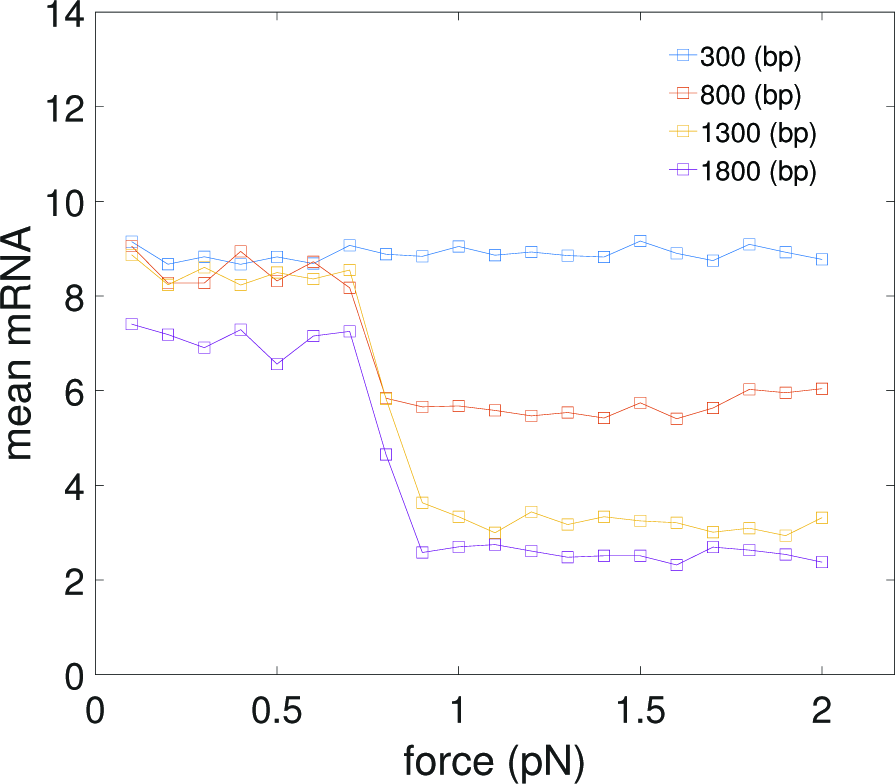
Effects of gene length on mRNA expression for an isolated gene transcribing against a single barrier as the external force is changed (composition details in SM). The remaining simulation parameters are the same as figure 3. Average mRNA expression for short genes is unaffected by changes in external force while long genes are repressed.

### Gene Expression Statistics

As is shown in fig. 5 the burst size for fixed mechanical properties is determined through the gene and barrier length. The gene specific stochastic rates controlling RNAP initiation *r* and degradation do not influence *m_c_* (shown in SM fig.S3). However, changing the nature of the mechanical properties of the drag associate with RNACs rotation changes *m_c_* as shown in SM fig.S3B. It is important to note though that while *m_c_* is a function only of the geometric properties of the gene (along with its intrinsic mechanical properties) the stochastic rates for RNAP initiation, mRNA degradation and relaxation have major effects on gene expression. In addition to setting time scales for the system these rates control characteristic times to frustration and relaxation, thus controlling how big a role mechanics and the burst size *m_c_* play in determining gene expression statistics. Thus the balance between frustration and relaxation are central to the quantitative understanding of transcription (figs.3 and 7). This effect offers a possible explanation for the absence of transcriptional bursting and its role in gene expression for some systems [32].

**FIG. 7:**
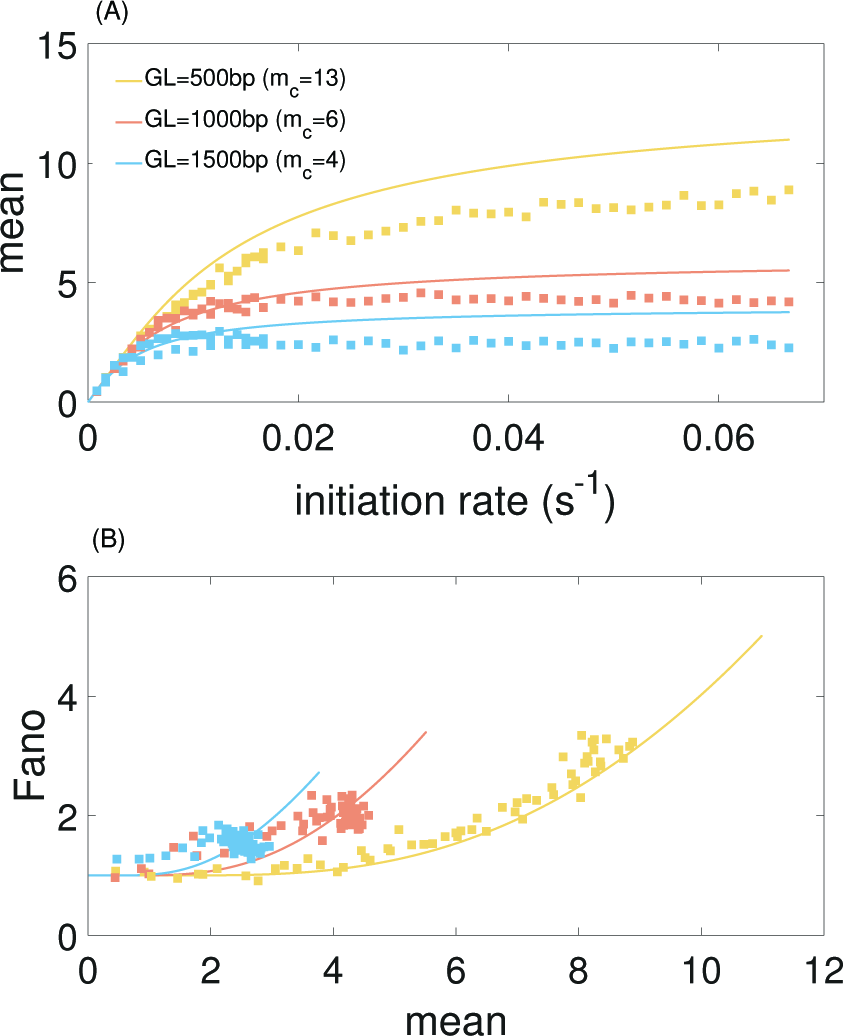
(A) Mean mRNA expression levels for genes with varying lengths GL (composition details in SM) for an isolated gene with mechanically interaction RNAP for increasing initiation rate. The remaining simulation parameters are the same as figure 3. The results show a clear limit to expression given by the burst size *m_c_*. (B) Relationship between mean expression levels and Fano factor for genes with varying lengths for an isolated gene with mechanically interaction RNAP showing a clear relationship between expression and noise as has been demonstrated experimentally.

In a previous work [32], we presented analytical calculations for the mean and variance of mRNA produced for a gene which had a hard cutoff to the number of transcripts *m_c_* which can be made without a relaxation event. It is therefore of interest to see how well that simple model can capture the consequences of bursting in this full mechanical treatment. Predictions for the mean mRNA levels and the associated Fano factor for this simple stochastic model are shown as solid curves for the corresponding parameter values in fig.7. The mean expression level is set by the burst size *m_c_* as

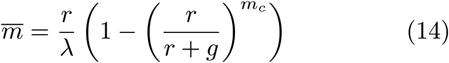

showing how the mechanical parameter *m_c_* controls the mean expression of a given gene. In general the expression for the Fano factor is complicated however for the system examined in fig. 7 with *λ* = *g* the Fano factor and mean can be written as a function of 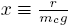

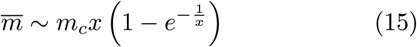

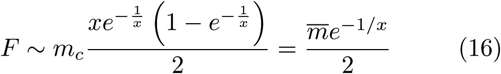

In general the analytical expression for the Fano factor is complicated [32].

Changing the characteristic time to frustration *m_c_/r* or the relaxation time 1/*g* changes the effects of the mechanical properties of transcription; for *m_c_/r < 1/g* the noise of the system is driven away from the Poissonian limit (Fano factor of one). As shown in fig. 7 the mechanical properties of transcription constrain the mean expression level and the noise. The mean expression rate in the fast initiation rate limit *r* → ∞ is set by the burst size *m_c_* as 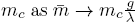 with a corresponding Fano factor of 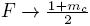. This interplay offers an explanation for both universal as well as gene specific aspects of transcriptional noise. Deviations between the analytical values and the simulated data can be attributed to a number of effects not considered in the simplified mathematical model such as RNAP occupancy exclusion. Incorporating additional regulatory effects such as repressors is not done here but was examined in our previous work [32].

These results clearly demonstrate the effects of the mechanical properties of transcription on the statistical properties of gene expression for isolated genes interacting with a mechanical barrier. However, we have not specified the nature of this barrier thus far. In the following section we consider the effects of a multiple-gene system on a shared piece of DNA where neighboring genes serve as twist barriers for each other.

### Interacting genes

In this section we will consider multiple genes acting in concert on a shared piece of DNA. In the same way that RNAPs on an isolated gene can interact, RNAPs on neighboring genes can also interact, mediated by DNA torsion. This will introduce a non-local correlation across genes. The ability for neighboring genes to influence each other has long been known [33] and a number of direct and indirect phenomena in gene expression have pointed to the location and orientation of genes as central determinants [9, 34] of their function. Additionally, a previous theoretical study [35] has examined interaction between genes due to SC controlled initiation. Here we will be concerned with how SC changes the ability of RNAC to elongate. As before, the motion of each RNAC will be governed by the DNA twist equation.

We will first examine the behavior of two genes. In addition to the role of gene length and barrier distance (now intergenic spacing), the role of gene orientation (which direction the RNAC travels) plays a pivotal role in determining the behavior of the respective genes. Two genes which are convergently or divergently oriented relative to one another will cause positive or negative SC to accumulate in the intergenic region, each serving as a twist barrier to the other. The accumulation of SC causes RNAC stalling and transcriptional bursting for genes which are convergently or divergently oriented in the same manner as an isolated gene with a static barrier. Consequentially, gene expression statics for such configurations are altered resulting in diminished expression levels as well generating correlations between genes as shown in fig.8. This effect is not present for tandemly oriented genes, in agreement with recent observations in synthetic systems [34]. The role of gene lengths and intergenic distances follows the results of isolated genes and is examined in the SM. Increasing the non-coding intergenic distances diminishes transcriptional bursting and allows for Pois-sonian gene expression.

**FIG. 8:**
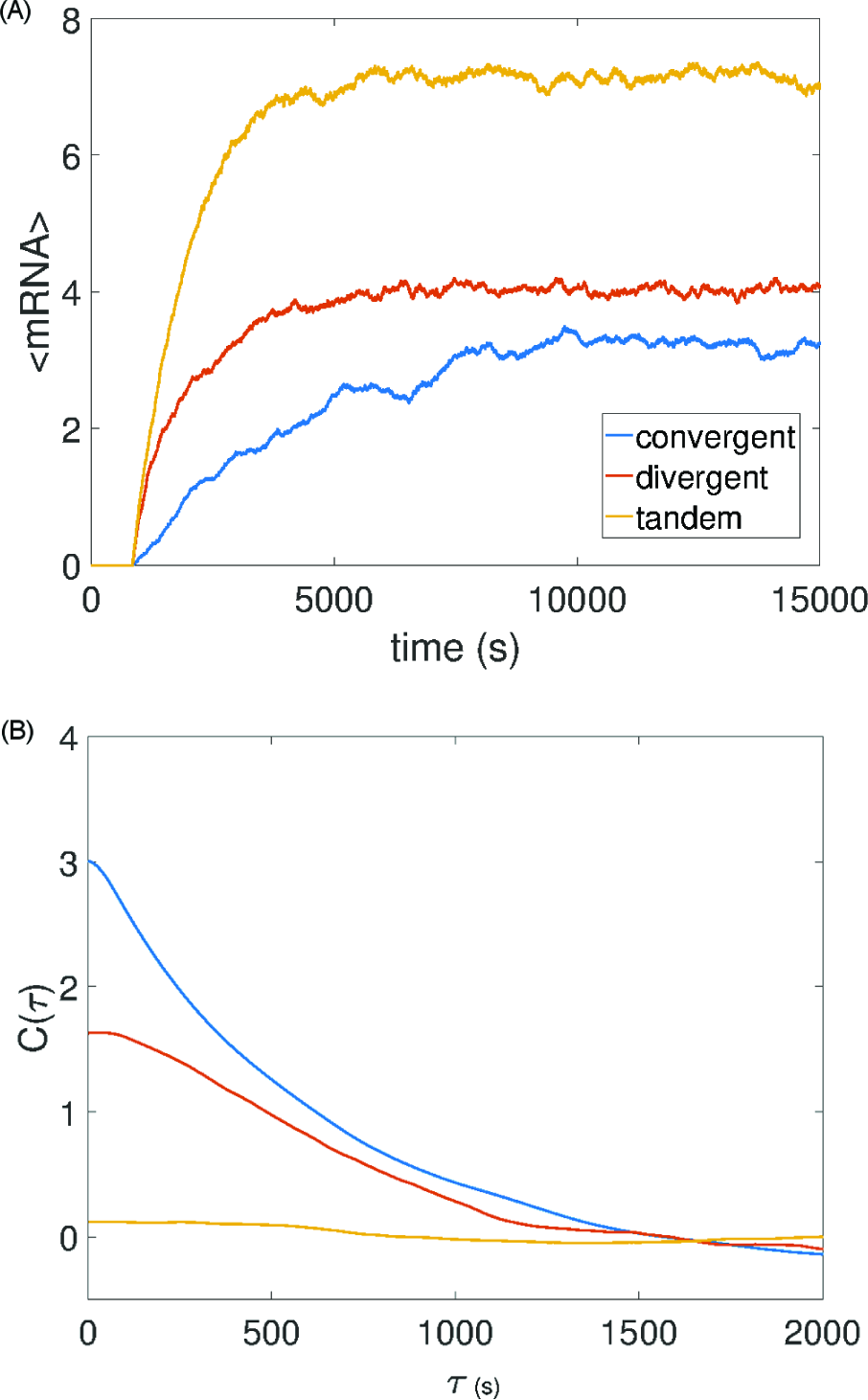
Effects of orientation on mRNA expression levels (A) and correlation (B). For genes which are convergently and divergently oriented SC builds up in the region between the genes causing correlations and altered gene expression (see SM for compositional details). (A) Average expression levels in time for many single trajectories showing a clear difference in mean expression levels as gene orientations are changed. (B) The mRNA correlation function between the two genes shows correlations change as a function of orientation. Gene and barrier lengths of 1 kbp and 10 kbp were used respectively. Correlation function computation are contained within the SM and the simulation parameters are the same as figure 3.

The non-local interaction between neighboring genes through DNA torsion and the effects of gene arrangement and intergenic length on gene expression offer an alternative method to alter gene expression beyond the ability of promoters and enhancers to interact [33, 36]. Additionally, these results offer an interesting role for non-coding DNA as a space for SC to accumulate, acting as a mechanical buffer for highly transcribed genes. This previously unconsidered phenomena will be important for the construction of synthetic genes and may serve as an explanation for many existing paradoxes in the arrangement of non-coding DNA in the genomes of living systems.

The behavior of small systems with multiple genes such as the one shown in figure 9A (composition details are in SM) follows from the behavior of two interacting genes. Many genes interacting in a shared region of DNA will affect each other in the same manner as we have previously outlined, where the lead or trailing RNAC serves as a twist barrier to the leading or trailing RNAC of the neighboring gene. Transcriptional bursting occurs for all the genes in a gene size and inter-gene length manner. Stochastic mRNA trajectories and statistics for the simple multiple gene system shown in figure 9A are shown in the SM. Changes in gene expression by tuning intrinsic parameters follow in the same manner as for isolated or pairs of genes.

**FIG. 9:**
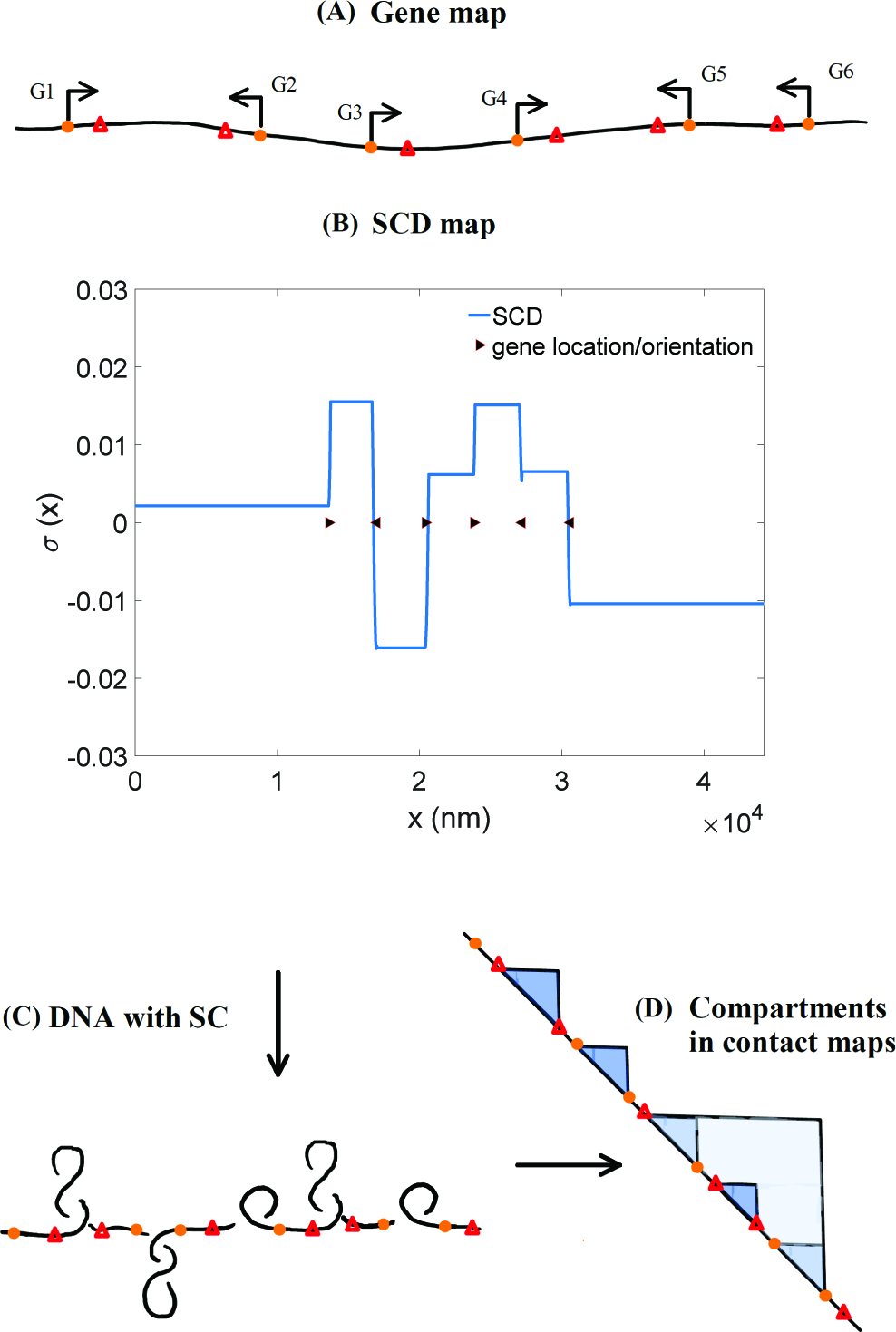
The geometric and compositional structure of genes shown in (A) determines the SCD structure of a region of DNA (B). Regions of SC influence the structural conformations realized by DNA (C) which will be manifested in chromosomal conformation contact maps (D) as domains. In some regions (such as for genes 3–6) the multiple levels of SCD can act collectively to create multiple levels of organization where smaller domains exist in larger domains.

### Super-coiling and DNA Structure

Recent observations have shown that regions of actively transcribed DNA have significant altered structural and epigenetic properties as compared to inactive regions [37]. These differences manifest themselves as chromatin domains of varying sizes in many organisms. The emergence of under and over-wound regions of inactive DNA flanked by torsionally neutral areas of active transcription as shown in figure 9 is a plausible explanation for the formation of chromatin domains which have been observed in structural studies of DNA in many organisms [38] and which in fact have been directly connected to active transcription in recent studies [39, 40]. The relative lengths, positions and orientations genes determine the regions of SC in DNA and will strongly influence the conformational structures realized by DNA in those regions. In a recent study in bacteria it has been shown that the longest most strongly transcribed genes serve as DNA domain boundaries [39]. Changing the location or lengths of these genes changed the boundary positions. Using the framework presented here a very similar behavior is observed where SCD domains move according to gene position, orientation and length (see fig.S1).

This connection offers an intriguing link between gene expression and DNA structure which has only been partially explored. Previous studies comparing SC regions to domains have used methods which detect DNA undertwisting [41]. These studies found partial agreement between SC regions and structural domains but this technique is not suited for observing fully plectonemic regions as should be expected in regions of significant SC. Additionally, increasing resolution in chromatin capture techniques have revealed previously missed small structural domains [40] and one must be careful to account for resolution limitations. Overlap between SC regions created through active transcription and domains is consistent with many known aspects of DNA structure and provides a compelling evolutionarily conserved mechanism for their formation in all organisms [40]. This mechanism can exist alongside additional mechanisms for alternating chromatin structure in higher organism, such as loop extrusion [42, 43]. Additionally changing the location and size of SC domains may lead to changes in histone occupancy resulting in altered epigenetic patterns and chromatin structure and mechanics.

As stressed throughout this work, the level of SC between genes is set by the mechanical limits to RNAP. This offers an additional constraint to understanding the impact of transcription on DNA structure which has already been tested in one study which clearly demonstrated the torques needed to induce the structural changes seen in chromatin studies to be within the capability of RNAP [44]. Indeed some coarse polymer simulations have already shown the ability of SC to generate a number of observed properties of chromatin structure [44, 45]. However additional study is still needed to fully understand the role of transcription in determining chromatin structure.

## DISCUSSION

The framework constructed here and the resulting phenomena offer a different perspective on the processes which determine and drive gene expression. Many results presented in this article make strong qualitative predictions which agree with experimental observations. Furthermore, the results of this article are able to explain the interaction of multiple genes on a shared DNA as well as offer an explanation for how external non-molecular changes can influence gene expression. These results are connected to the role of DNA structure and mechanics in transcription. The non-local interaction and the resulting structural changes to DNA can connect disparate levels of cellular structure and function.

Examining the mechanical properties of transcription from this perspective can tie together many disparate phenomena in biology. The non-local interaction and the resulting structural changes to DNA which effect genes on the same piece of DNA generate a number of interesting phenomena. As shown in the paper this includes transcriptional bursting (fig 3) and its origins as a geometric property of a gene and it’s surroundings (fig 5). The geometric origin of bursting allows us to make predictions on the burst size *m_c_*. The relationship between burst size and gene expression statistics is readily observed and a plausible explanation for the observed generic relationship between gene expression levels and noise is given. The non-local interaction between genes leads to significant changes in gene expression, not do to the role of transcription factors, but do to the behavior of the surrounding genes. Additionally this phenomena offers an interesting role for non-coding regions of DNA as a mechanical buffer to active transcription of surrounding genes. Due to the strong ability of RNAP to introduce torsion into DNA, transcription introduces significant changes in the SC levels throughout DNA resulting in over and under winding in non-active areas and neutral SC in active regions of transcription. This phenomena offers a compelling mechanism for the formation of active and inactive domains observed in structural studies of DNA across all organisms.

Throughout the article a simple method for understanding the torsional response of DNA (eq. 5) was utilized. As explained above, this formulation relies on a separation of time-scales between DNA mechanics and RNAC movement allowing for the use of an equilibrium formulation of DNA mechanics which assumes an instantaneous torsional response of DNA. While this method is fine for relatively short pieces of DNA, simulating long intergenic regions might require the use of more sophisticated models of DNA which include torsion transport and explicitly incorporate the interplay between twist and writhe. Introduction of DNA proteins such as histones could also be accomplished (changing the torsional response of DNA) though we do not believe the qualitative predictions made in the article will be changed. Incorporating more sophisticated models of DNA into the framework used here can be done without any obvious barriers and is left for future work.

Additional processes beyond RNAP velocity are effected by SC. One obvious process is initiation, which relies on DNA unwinding to operate. Incorporating this phenomena into the model would amount to making stochastic initiation rate SC dependent and some theoretical work has been completed examining it’s effect in gene expression [46]. Additionally active reorganization of DNA due to transcription introduces a mechanism by which transcription increases access to enhancers and repressors. While not considered here these effects are interesting and potentially important to understanding transcription in living systems. Future work will can be done to introduce these processes and others into the model presented.

Finally, the theoretical discussion within this article has not centered on any particular organism. Though we have neglected a number of potentially important organism specific effects, we believe that the presented framework is capable of capturing the same phenomena in many organisms. As stated in the introduction, we have been concerned with the conceptual, broad characterization of the mechanical properties of transcription and their role in gene expression and DNA organization.

In conclusion it has become clear over the past several decades that chemical and physical processes play a central role in determining where and when a particular gene is active. In this work we have only laid out some of the most basic features of the physical act of transcription for multiple RNAPs acting in a multi-gene system. Examining the physical side to transcription has uncovered a role for genome composition (gene orientation, size, and intergenic distance) beyond the organization of regulatory elements. Though many aspects of the model are simple, additional theoretical and experimental woks can constrain and refine the models precision and offer increasing levels of prediction. Future efforts to understand the precise mechanisms of cellular function will have to take the effects outlined here under consideration.

## ACKNOWLEDGEMENTS

This work was supported by the National Science Foundation Center for Theoretical Biological Physics (Grant NSF PHY-1427654).

## SUPPLEMENTARY MATERIAL

### Boundary Conditions

In the framework constructed in the RNACs at the ends of genes interact with the closest RNACs on neighboring genes to result in a complete set of LNCs 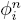 for all RNACs at positions 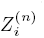. These of course assume that there is at least one RNAC at the neighboring gene; if not we just skip genes until the next RNAC is found.For this to work we must have conditions for all possible gene configurations. We will label genes with positive + and the opposition genes −. Thus for any given gene there are 8 possible orientations given neighbors on each side (±, ±, ±). Explicitly We have

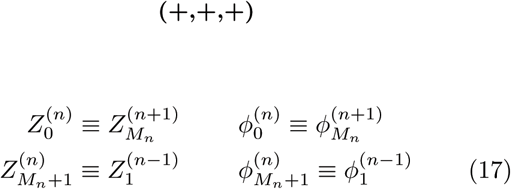

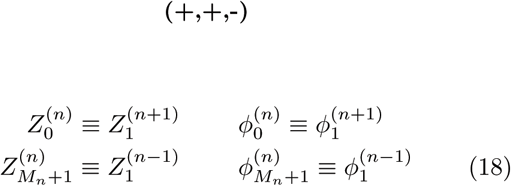

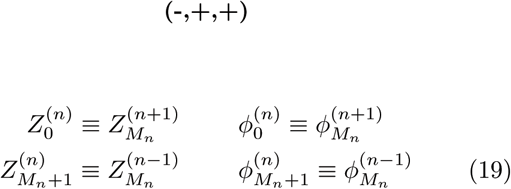

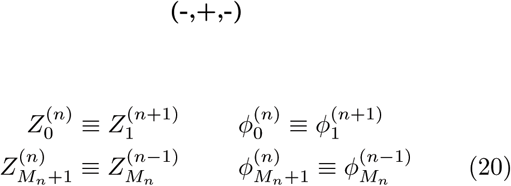

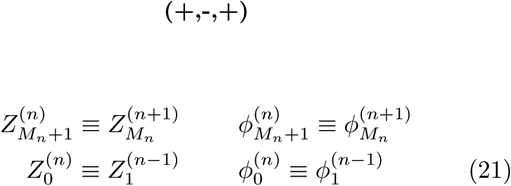

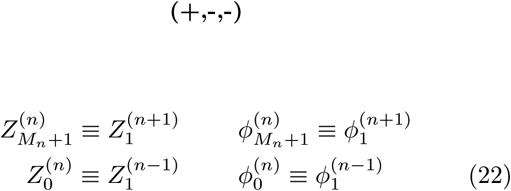

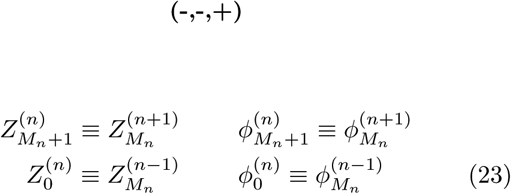

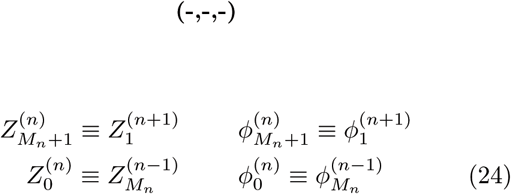

If the left or right side of the gene contains no additional genes the boundary conditions are set to have either free (torsionally unconstrained) motion by adding an additional LNC matching the end RNACs LNC or fixed (torsionally constrained) motion by an additional static LNC.

### Correlation Function

The correlation function shown in fig. 8 was calculated using the mRNA content in time of the left *m_L_*(*t*) and right genes *m_R_(t)* with different orientations using the formula

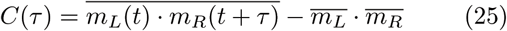

**FIG. S1:**
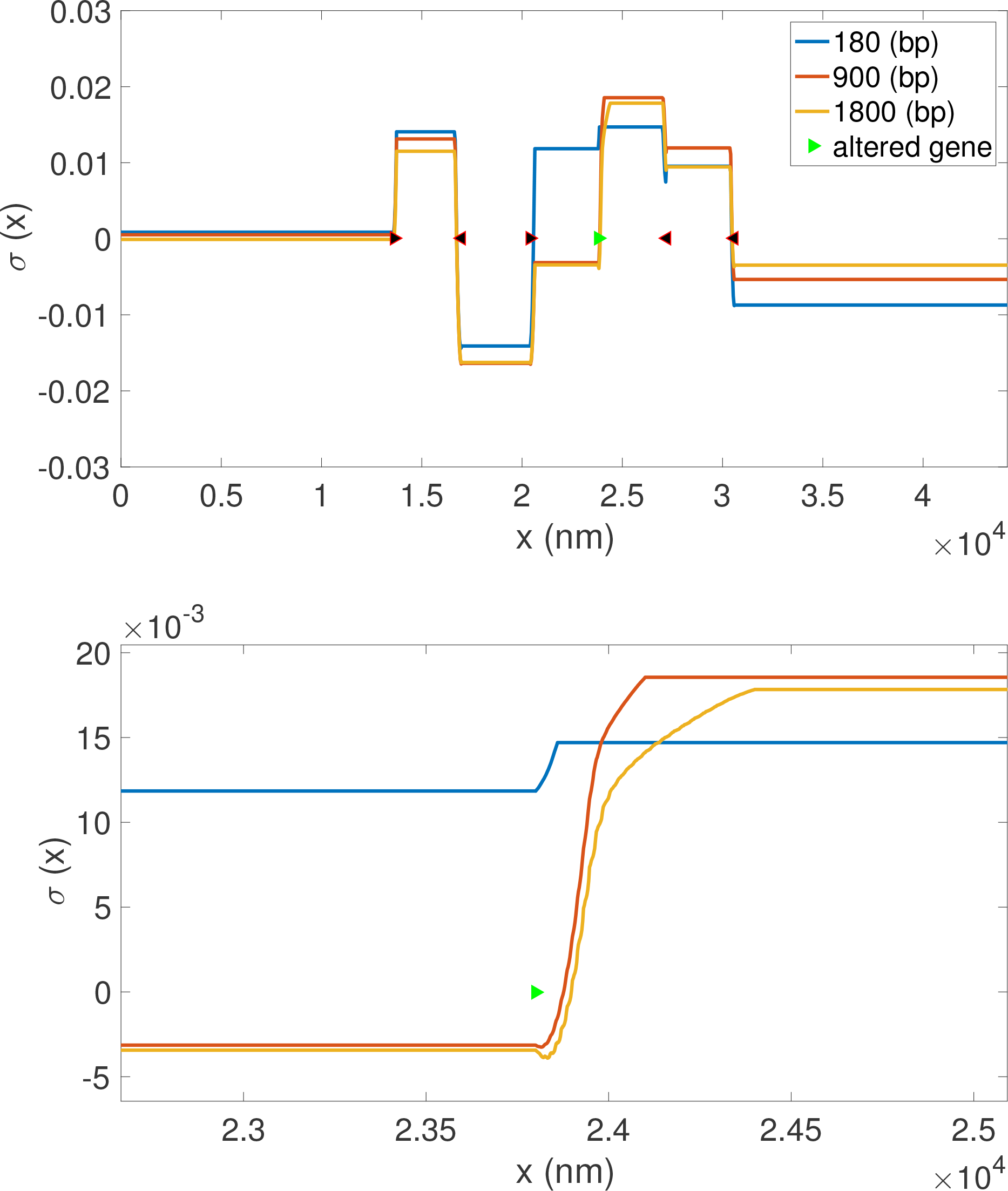
The geometric and compositional structure of the multi-gene system shown in fig.9A (with compositional details in fig.) determines the SC structure of a region of DNA (see fig.9B). Changing the length of a particular gene leads to SC structure rearrangement and thus DNA conformational changes. This can be see by changing the length of one particular gene in a multi-gene system at the system level (A) and in detail over the changed detail in (B).

**FIG. S2:**
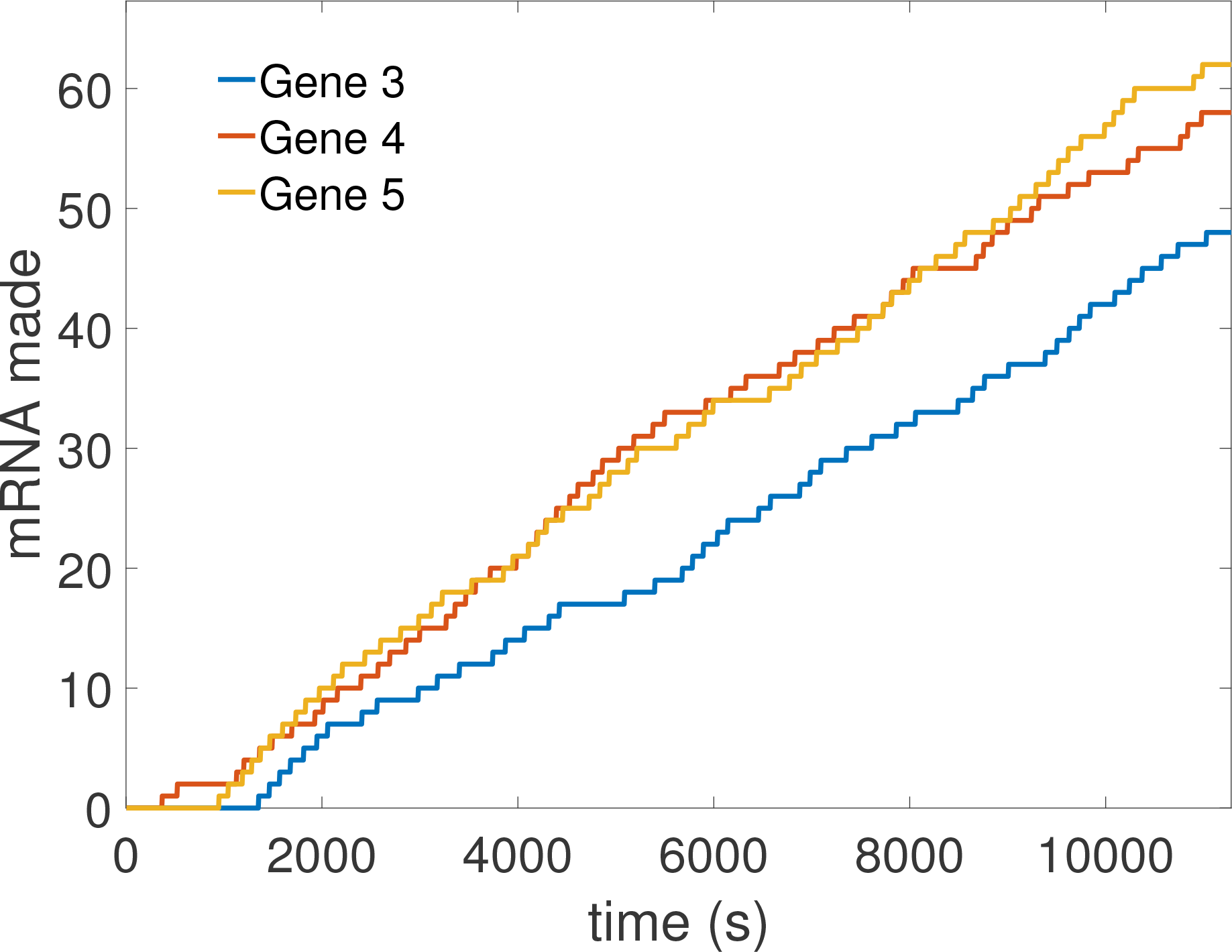
Bursts in production of mRNA shown for three of the genes of the system shown in figs.9, S1.

**FIG. S3:**
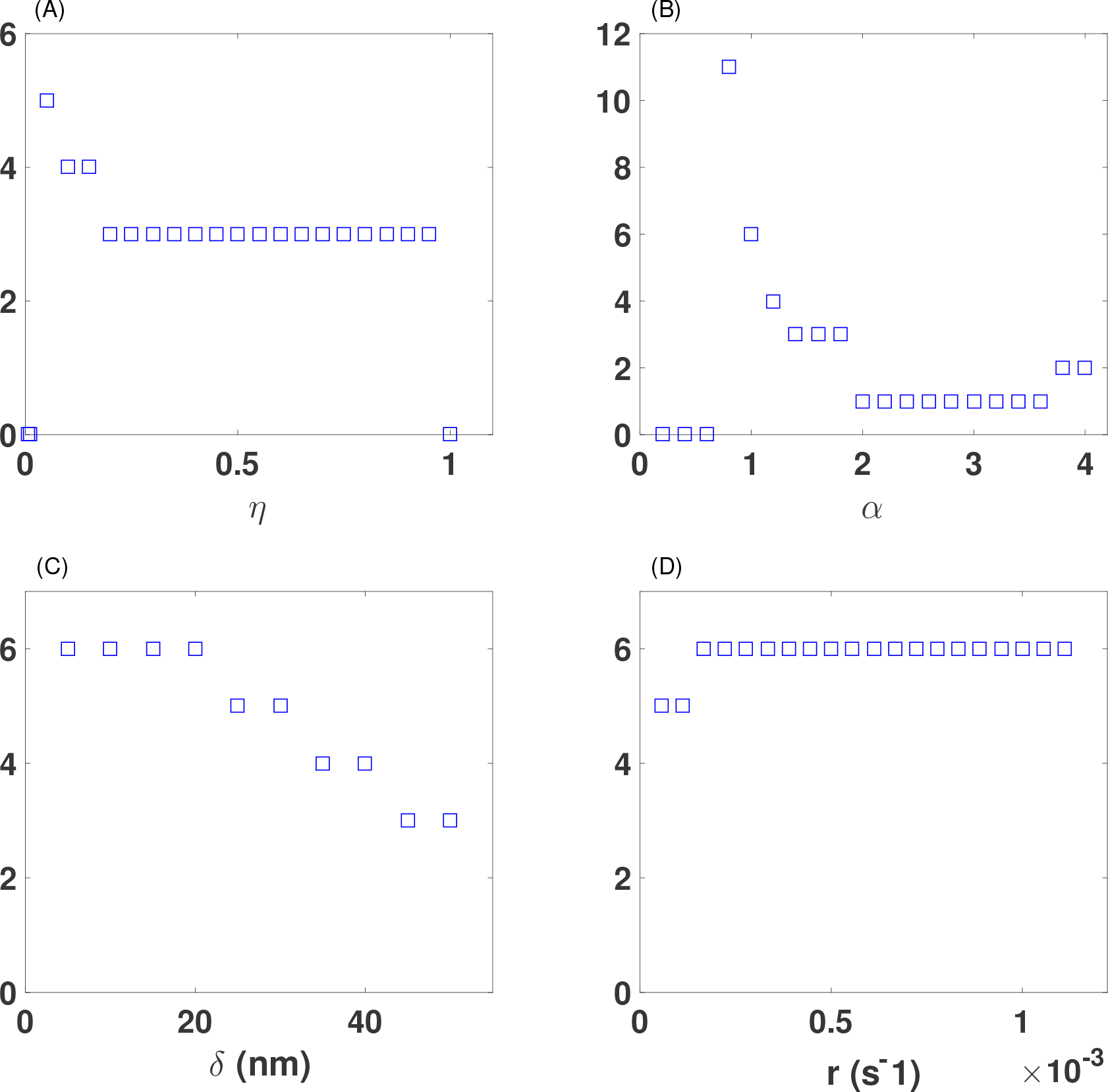
Burst size *m_c_* dependence on drag coefficient *η*,drag exponent **, RNAC spacing *δ*, and initiation rate *r* for an isolated gene with mechanically interacting RNAP. Gene and barrier lengths are 1kbp and 10 kbp respectively. Unless being varied the simulation parameters are {*r, λ, g*} = {2^−1^,1/20,1/20(*min*^−1^)} were used. The mechanical parameters are {*v_0_, η, α, χ, f*} = {20, 1/20, 1, 10, 1}

**FIG. S4:**
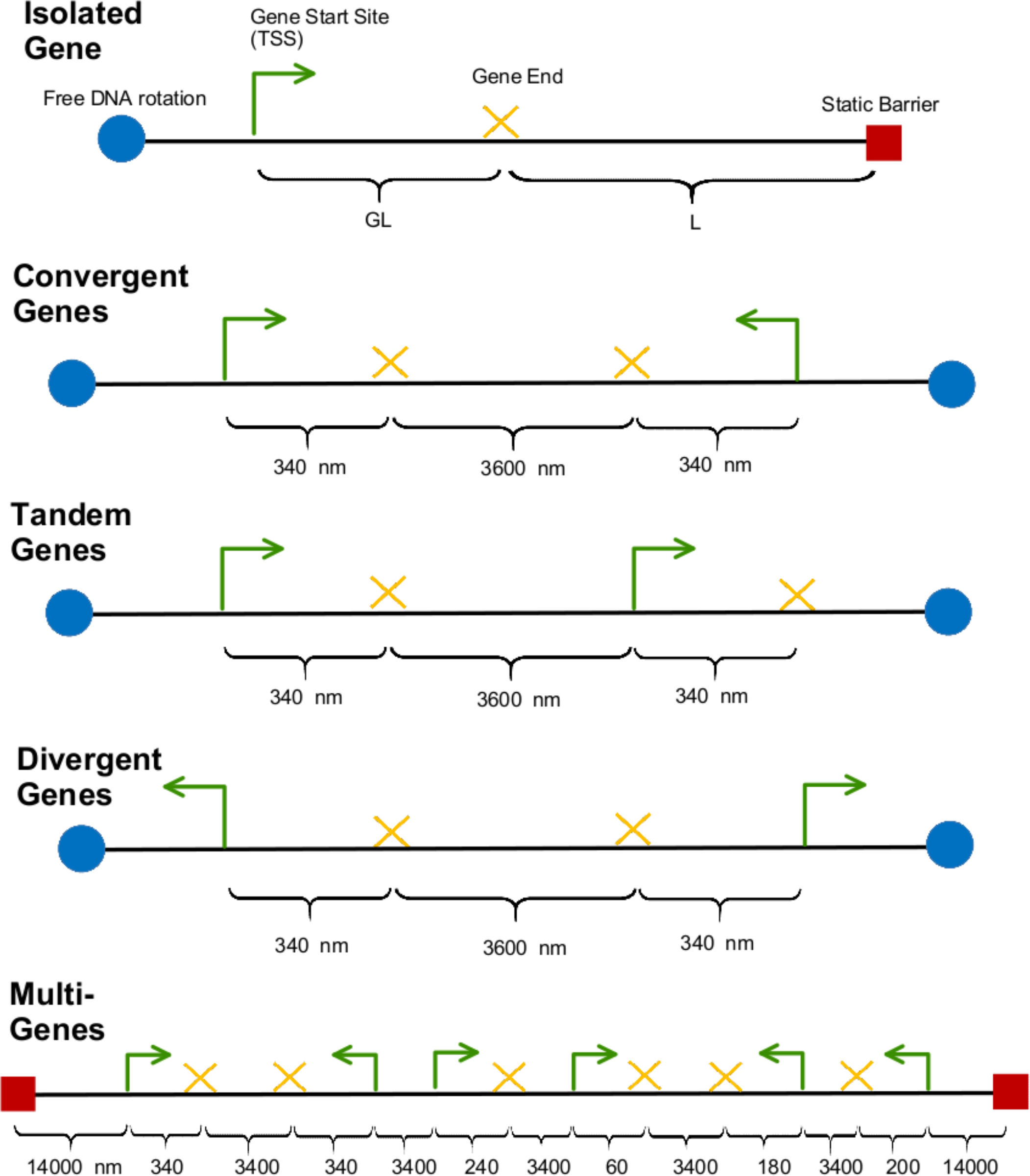
Compositional details for genetic systems used in simulations.

